# Adolescent Thalamocortical Inhibition Alters Prefrontal Excitation-Inhibition Balance

**DOI:** 10.1101/2023.11.22.568048

**Authors:** David Petersen, Ricardo Raudales, Ariadna Kim Silva, Christoph Kellendonk, Sarah Canetta

**Author notes:** Equal contribution (Christoph Kellendonk, Sarah Canetta).

## Abstract

Adolescent inhibition of thalamo-cortical projections from postnatal day P20-50 leads to long lasting deficits in prefrontal cortex function and cognition in the adult mouse. While this suggests a role of thalamic activity in prefrontal cortex maturation, it is unclear how inhibition of these projections affects prefrontal circuit connectivity during adolescence. Here, we used chemogenetic tools to inhibit thalamo-prefrontal projections in the mouse from P20-35 and measured synaptic inputs to prefrontal pyramidal neurons by layer (either II/III or V/VI) and projection target twenty-four hours later using slice physiology. We found a decrease in the frequency of excitatory and inhibitory currents in layer II/III nucleus accumbens (NAc) and layer V/VI medio-dorsal thalamus projecting neurons while layer V/VI NAc-projecting neurons showed an increase in the amplitude of excitatory and inhibitory currents. Regarding cortical projections, the frequency of inhibitory but not excitatory currents was enhanced in contralateral mPFC-projecting neurons. Notably, despite these complex changes in individual levels of excitation and inhibition, the overall balance between excitation and inhibition in each cell was only changed in the contralateral mPFC projections. This finding suggests homeostatic regulation occurs within subcortically but not intracortical callosally-projecting neurons. Increased inhibition of intra-prefrontal connectivity may therefore be particularly important for prefrontal cortex circuit maturation. Finally, we observed cognitive deficits in the adult mouse using this narrowed window of thalamocortical inhibition (P20-P35).

**Significance Statement:** Connectivity between two brain regions, the thalamus and the prefrontal cortex, has been found to be reduced in patients with schizophrenia. Neuronal activity in thalamo-cortical projections is important for the proper development of sensory cortices. How thalamo-cortical activity regulates prefrontal cortex development is less well understood. Here, we show that decreasing activity in thalamo-prefrontal projections in mice during early adolescence alters synaptic connectivity to distinct neuronal projections within the prefrontal cortex that are already evident in adolescence. While some of these changes can be explained by reduced thalamo-cortical projections, other adaptations are intrinsic to the prefrontal cortex. These findings implicate adolescence as a critical period of cortical development and demonstrate this period as a potential target for therapeutic intervention.

## Introduction

The prefrontal cortex (PFC) supports high level cognitive functioning including working memory and other executive functions (Fuster, 2001; Miller et al., 2018). An impairment in the development or maturation of the prefrontal cortex during adolescence has been postulated to contribute to the cognitive deficits observed in patients with schizophrenia (Feinberg, 1982; Paus et al., 2008; Gogtay et al., 2011; Hoops and Flores, 2017; Larsen and Luna, 2018; Sakurai and Gamo, 2019; Chini and Hanganu-Opatz, 2021; Dienel et al., 2022; Yang and Tseng, 2022). Thalamic inputs regulate sensory cortical maturation during sensitive time windows of development (Wiesel and Hubel, 1963; Nabel and Morishita, 2013). We recently established in the mouse that thalamic input activity during adolescence is necessary for adult prefrontal cortex function and behavior (Benoit et al., 2022). More specifically, we found that decreasing activity in the mediodorsal thalamus from postnatal day 20-50 using pharmacogenetic tools led to deficits in the performance of two prefrontal dependent cognitive tasks addressing working memory and attentional set shifting. These deficits were measured in the adult mouse and associated with decreased anatomical thalamocortical projections and decreased functional excitatory post-synaptic currents measured in layer 2/3 prefrontal cortical pyramidal cells. Moreover, mice with thalamic inhibition during adolescence showed decreased correlated activity between neurons in the medial prefrontal cortex (mPFC) while performing an attentional set shifting task and the ability of the prefrontal cortex to decode task outcomes was impaired (Benoit et al., 2022). Notably, decreasing the activity of thalamo-prefrontal projections for a comparable length period in adulthood did not lead to the same long-lasting effects.

These findings established that thalamo-prefrontal activity is required during adolescence to sustain thalamocortical connectivity, prefrontal network function and cognitive behavior in adulthood. However, it was not clear whether these effects on circuit connectivity were apparent immediately after the adolescent manipulation or if they require maturation into adulthood to manifest. Moreover, it was not clear whether the effects of decreased adolescent thalamo-prefrontal activity were restricted to the reduction of excitatory thalamic afferents or whether this manipulation led to more widespread changes in the intrinsic connectivity of the prefrontal cortex.

To address these questions, we undertook a detailed study of prefrontal cortical connectivity using slice electrophysiology twenty-four hours following pharmacogenetic inhibition of thalamo-prefrontal projections from postnatal day P20-35. We chose this reduced window of inhibition because prior work demonstrated changes in the density of thalamic projections to the mPFC following adolescent thalamic inhibition were already present at this time (Benoit et al., 2022). We first recorded excitatory and inhibitory currents from layer II/III neurons as previously done in the adult (Benoit et al., 2022) to establish whether the observed deficit in excitation arises during adolescence. We then restricted our recordings to specific mPFC projection neurons identified by their projection target and cortical layer. We found excitatory post synaptic currents are reduced in layer II/III cells during adolescence, as observed in adult mice. When analyzing specific projections, we found a reduction in frequency of excitatory and inhibitory currents to layer V/VI cortico-thalamic cells and layer II/III cortico-accumbens-projecting cells while layer V/VI cortico-accumbens-projecting neurons displayed increased amplitudes of excitatory and inhibitory currents. While we observed changes in excitation or inhibition in subcortically-projecting cells, the excitation/inhibition balance was not affected, suggesting homeostatic mechanisms may offset decreased excitatory inputs. In contrast, callosal cortico-cortical projection neurons displayed a significant decrease in E/I balance. Together, these data show a complex adaptation of cortical networks in response to decreased thalamo-cortical inputs during adolescence and suggest that activity in callosally-projecting cells in mPFC may be most profoundly affected due to a lack of homeostatic adaptation.

To determine whether this narrowed window (P20-P35) of adolescent thalamocortical inhibition would impact cognitive flexibility in the adult mouse, we performed an attentional set shifting behavioral task. Like P20-50 inhibition (Benoit et al., 2022) inhibition from P20-35 was sufficient to induced long lasting deficits in cognition.

## Materials and Methods

### Animal Husbandry

All procedures were performed in accordance with guidelines approved by the Institutional Animal Care and Use Committees at Columbia University and the New York State Psychiatric Institute (protocol NYSPI 1589). Animals were housed under a 12-h light/12-h dark cycle in a temperature-controlled environment (22 °C, humidity of 30–70%) with food and water available ad libitum, unless otherwise noted. C57/ BL6 females and males (Jackson Laboratories, 000664) were used for all experiments. Littermates were randomly assigned to each experimental group, with random distribution across males and females. Mice were housed together with dams and littermates. Offspring were weaned at P28, and group housed with same-sex littermates (no more than five mice per cage). For thalamic inhibition, mice were given intraperitoneal (i.p.) injections of JHU37160 (J60, a CNO analog) dissolved in 0.9% sterile saline at 0.1 mg/kg twice per day (Bonaventura et al., 2019). J60 was used as an alternative to the widely used clozapine-N-oxide (CNO) because J60 is known to have less off-target effects. New evidence has shown CNO might be reverse metabolized into clozapine, a common antipsychotic medication with high affinity to dopamine D2 and serotonin 2A receptors (Manvich et al., 2018). All mice were given J60 regardless of viral vector, except those given saline with hM4D expression. Surgeries were conducted at P12-13, and all mice were injected from P20 to P35. For electrophysiological experiments, 24 hours after the final J60 injection, mice were anesthetized with isoflurane and decapitated for slice electrophysiology at P35 (+/- 4 days). For behavioral experiments, mice were tested in an attentional set shifting (ASST) task beginning in adulthood (>P90). Throughout data collection and analysis, experimenters were blinded to the group of the animal. We based sample sizes on previous experiments, and no statistical methods were used to calculate sample sizes.

### Surgical Procedures

For viral injections at P13, mice were anesthetized with isoflurane and head-fixed in a stereotactic apparatus (Kopf). Mice were injected bilaterally in the midline thalamus with AAV5-hSyn-DIO-hM4D-mCherry (Addgene, 44362) or a control virus, AAV5-hSyn-DIO-mCherry (Addgene, 50459), at a volume of 0.25 µl (0.1µl/min). Mice were also injected bilaterally in the mPFC with retrograde AAV-hSyn-Cre-WPRE-hGH (Addgene, 105553) at a volume of 0.2 µl (0.1 µl/min). For the tracer experiments, mice were injected with retrograde AAV-hSyn-EGFP (Addgene, 50465) at a volume of 0.25 µl for the MD, 0.2 µl for the mPFC, and 0.1µl for the NAc. The following were the P13 coordinates: MD thalamus, −0.85 anterior–posterior (AP), ±0.20 medial–lateral (ML) and −3.25 dorsal–ventral (DV, zero at brain surface); mPFC, 2.4 AP, ±0.2 ML and −1.9 DV (zero at brain surface); NAc, 1.7 AP, ±1.1 ML and −5.07 DV. All coordinates were zeroed at bregma unless otherwise noted. Histology images were collected from each animal for each cohort, with at least 6 slices taken for each animal to determine viral spread and confirm targeting.

### Slice Electrophysiology

Whole-cell voltage clamp recordings were performed in layer II/III mPFC pyramidal cells and layer V/VI mPFC pyramidal cells. Recordings were obtained with a Multiclamp 700B amplifier (Molecular Devices) and digitized using a Digidata 1440B acquisition system (Molecular Devices) with Clampex 10 (Molecular Devices) and analyzed with pClamp 10 (Molecular Devices). Following decapitation, 300-µm slices containing mPFC were incubated in artificial cerebral spinal fluid containing 126mM NaCl, 2.5mM KCl, 2.0mM MgCl2, 1.25mM NaH2PO4, 2.0mM CaCl2, 26.2mM NaHCO3 and 10.0mM d-glucose (pH 7.45, 310 mOsm) bubbled with oxygen at 32 °C for 30min before being returned to room temperature for at least 1 hour before use. During recording, slices were perfused in room temperature artificial cerebral spinal fluid at a rate of 5 ml/min. Electrodes were pulled from 1.5-mm borosilicate glass pipettes on a P-97 puller (Sutter Instruments). Electrode resistance was typically 1.5–3MΩ when filled with internal solution consisting of 130mM cesium hydroxide monohydrate, 130mM D-gluconic acid, 10mM HEPES, 2.0mM MgCL2, 0.2mM EGTA, 2.5mM Mg-ATP, 0.3mM Na-GTP, and 5mM Lidocaine N-ethyl bromide (pH 7.3, 277mOsm).

#### mPFC recordings

Animals were sacrificed for recordings at P35 after the adolescent manipulation. mPFC pyramidal cells were visually identified based on their shape and prominent apical dendrite at ×40 magnification under infrared and diffusion interference contrast microscopy. Recordings were taken from either layer II/III of the mPFC (PrL) or layer V/VI. Layer II/III was defined as any cell lying 150µm-350µm from the pial surface. Layer V/VI was defined as cells within 400µm-600µm from the pial surface. Spontaneous excitatory post synaptic currents (sEPSCs) were recorded in voltage clamp at a holding potential of −65mV, and spontaneous inhibitory post-synaptic currents (sIPSCs) were recorded in voltage clamp at a holding potential of +10mV. The combination of the intracellular solution and the selected holding potentials allowed us to isolate EPSCs and IPSCs from one another without the use of blockers, which we verified by demonstrating that sEPSCs were completed blocked by administration of 50 uM APV ((2R)-amino-5-phosphonovaleric acid) and 20 uM CNQX (6-cyano-7-nitroquinoxaline-2,3-dione), but not 20 uM bicuculine, and vice versa for sIPSCs.60 seconds of the current recording for each condition were analyzed. Recordings were filtered with a 2 kHz eight-pole low-pass Bessel filter, and sEPSCs and sIPSCs were detected using MiniAnalysis (Synaptosoft). Frequency, amplitude, and instantaneous frequency were calculated for all traces. Frequency was calculated by dividing the total number of events by time (60s). Amplitude was the distance from baseline to the tip of the event. Instantaneous frequency was the inverse of the interevent-interval converted to a frequency. All event data were averaged by cell.

### Attentional Set Shifting Behavioral Task

At P90, mice were gradually restricted to 85% of their body weight. Mice were habituated to the testing arena on day 1. On days 2–3, they were trained to dig in both bedding media (corn cob and paper pellet, both unscented) to obtain a food reward. Once mice dug reliably, testing began. For each trial, mice were placed at the opposite end from two terra cotta bowls containing different odor/medium combinations. For IA, mice needed to learn that the cinnamon scent, not the paprika scent, predicted a Honey Nut Cheerio reward, irrespective of the bedding medium. For the first five trials, mice could explore both bowls until they found the reward, but the trial was only scored as correct if the animal initially chose the correct bowl. From the sixth trial onward, once the mouse began digging in a bowl, the entrance to the other bowl was closed off. The criterion was reached when the mouse made eight of ten consecutive correct choices. If the mouse did not meet the criterion in 30 trials, the animal did not advance to the next stage. If the mouse did reach the criterion, the extra-dimensional set shifting (EDSS) portion of the task began. In EDSS, the animal needed to learn that the type of bedding medium (paper pellets, not corn cobs) predicted the Honey Nut Cheerio reward irrespective of odor. The criterion was reached when the mouse made eight of ten consecutive correct choices. All behavioral tasks were conducted during the light cycle.

### Statistics

Statistical analysis and graph preparations were done using Prism 9 software (GraphPad Software), JASP (JASP Team (2023), JASP (Version 0.17.3)) or custom scripts in R. Linear mixed effects models with individual animal used as a random-effects grouping factor were used to analyze slice physiology. Linear mixed effects models were used over a common ANOVA or t-test to account for the possible non-independence of measurements from cells from the same animal. Prior to analysis, all datasets were tested for outliers, and they were removed using the following equation (Q1-1.5 x IQR or Q3 + 1.5 x IQR), which identifies datapoints +/- 2.698σ from the mean. For the nonspecific L II/III experiment, 2 outliers were removed from the EPSC hM4D+Saline amplitude group, 1 outlier was removed from the IPSC hM4D frequency group, and 1 outlier was removed from the IPSC hM4D instantaneous frequency group. For the cortico-accumbens experiment, 2 outliers were removed from the IPSC hM4D group. For the cortico-cortical population experiment, 2 outliers were removed from the IPSC mCherry frequency group. For the E/I balance analysis, 2 outliers were removed from the hM4D cortico-cortical frequency group. An independent samples t-test was used to analyze behavior.

## Results

### Adolescent thalamocortical activity regulates excitatory transmission to superficial mPFC

We first performed patch clamp recordings in brain slices from prelimbic layer II/III neurons, which is the mPFC population of projection neurons that obtain the most prominent input from the MD (Anastasiades and Carter, 2021). To inhibit thalamic activity during adolescence on postnatal day 13 (P13), we injected an adeno-associated virus (AAV) carrying a Cre-dependent version of the inhibitory designer receptor hM4D into the thalamus of C57 mice and a retrograde virus carrying Cre recombinase, rgAAV-CRE, into the mPFC. Retrograde transport will cause Cre recombinase expression in the MD, leading to hM4D expression selectively in medial prefrontal-projecting neurons (Fig. 1a). We then chronically inhibited these cortically-projecting MD cells with the hM4D ligand JHU37160 (J60) (0.1 mg/kg) from postnatal day 20 until the day before they were recorded (P35±4days) (Fig. 1b). To control for the possible non-ligand-dependent effects of the hM4D receptor, we also included a group of mice expressing hM4D that were injected with saline during the same developmental window. Twenty-four hours after the last J60 injection (at P35±4days), we recorded layer II/III neurons and analyzed excitatory and inhibitory post-synaptic currents.

**Figure 1.**
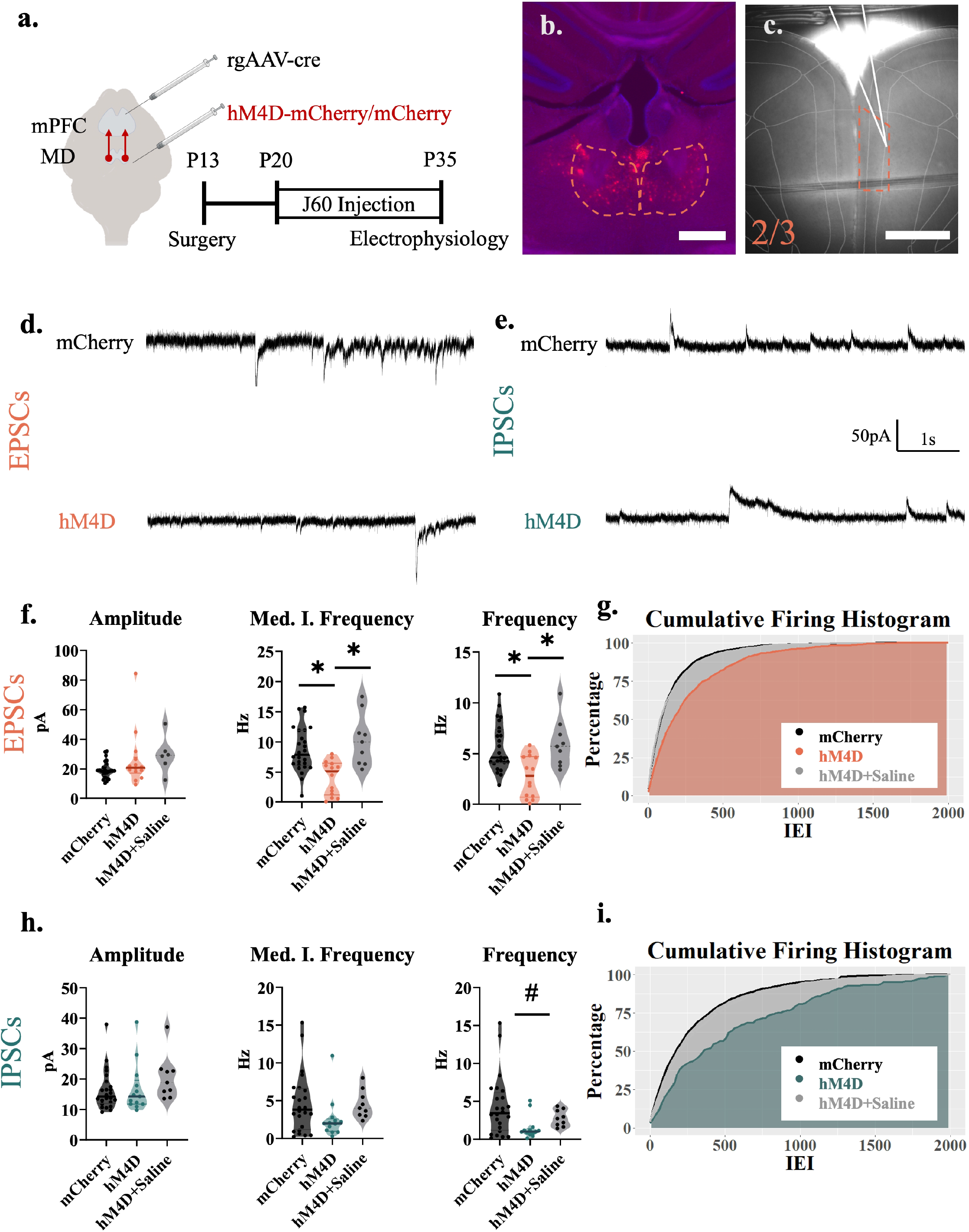
Adolescent thalamocortical activity regulates excitatory transmission to superficial mPFC. **a**, Adolescent experimental timeline, and schematic. Whole-cell patch clamp recordings were made from pyramidal cells in layer II/III of the mPFC from hM4D and control mice. These pyramidal cells receive excitatory inputs from the thalamus as well as inhibitory inputs from local interneurons; created with https://biorender.com. **b**, Example image illustrating hM4D-mCherry expression in the mediodorsal thalamus (dotted outline) in adolescent animals. Scale bar represents 300µm. **c**, Example image illustrating the recording location in the prelimbic (PrL) subregion (orange dotted outline) of the medial prefrontal cortex. All recordings for this experiment were conducted in this region approximately 200-350µm from the pial surface (L II/III) denoted by the orange dotted line. Recording pipette outlined in white. Scale bar represents 500µm **d**, Representative traces showing sEPSCs of hM4D animals and mCherry controls. **e**, Representative traces showing sIPSCs of hM4D animals and mCherry controls. **f**, sEPSC frequency (right) and median instantaneous frequency (middle) is significantly reduced following adolescent thalamic inhibition relative to control mice, but sEPSC amplitude (left) is unchanged; mCherry control, *n* = 26 cells and 6 animals; hM4D, *n* = 14 cells and 5 animals; hM4D+Saline *n*=9 cells and 2 animals; frequency: mCherry control 4.617(3.050) Hz, hM4D 2.833(3.858) Hz, and hM4D+Saline 5.733(2.700) Hz, contrasts of estimated marginal means revealed a significant difference between hM4D animals and both mCherry, *z* = - 3.311.129, d.f. = ∞ \**P_bonferroni_* = 0.003, and hM4D+Saline, *z* = 3.035, d.f. = ∞ \**P_bonferroni_* = 0.007 but no significant difference between mCherry and hM4D+Saline, *z* = 0.501, d.f. = ∞ \**P_bonferroni_* = 1.000; median instantaneous frequency: mCherry control 7.892(5.340) Hz, hM4D 5.126(4.882) Hz, hM4D+Saline 10.005(4.989) Hz; contrasts of estimated marginal means revealed a significant difference between hM4D animals and both mCherry, *z* = −3.032, d.f. = ∞ \**P_bonferroni_* = 0.007, and hM4D+Saline, *z* = 3.348, d.f. = ∞ \**P_bonferroni_* = 0.002 but no significant difference between mCherry and hM4D+Saline, *z* = 1.097, d.f. = ∞ \**P_bonferroni_* = 0.818; amplitude: mCherry control 18.785(5.250) pA, hM4D 20.916(4.882) pA, and hM4D+Saline 28.786(5.782) pA. All reported values are Median(Interquartile Range). **g**, Graph of the cumulative firing distribution for sEPSCs in L 2/3 pyramidal cells in the mPFC, IEI represents the interevent-interval between spontaneous events. **h**, sIPSC frequency (right) may trend downwards but median instantaneous frequency (middle) and amplitude (left) are unchanged following adolescent thalamic inhibition relative to control mice; mCherry control, *n* = 26 cells and 6 animals; hM4D, *n* = 14 cells and 5 animals; frequency: mCherry control 2.500(3.267) Hz, hM4D 1.000(0.550) Hz and hM4D+Saline 2.717(1.833); median instantaneous frequency: mCherry control 3.826(5.039) Hz, hM4D 1.989(1.256) Hz, and hM4D+Saline 4.014(2.366; amplitude: mCherry control 14.264(5.638) pA, hM4D 14.3(6.075) pA, and hM4D+Saline 18.849(6.737). All reported values are Median(Interquartile Range). **i**, Graph of the cumulative firing distribution for sIPSCs in L 2/3 pyramidal cells in the mPFC, *P<0.05, #P∼0.10.

Using linear mixed effects models, we found that inhibition of thalamo-cortical projections from P20-35 decreased the frequency [*F*(2, 5.93) = 6.854, *p* = 0.029] and spatial distribution (instantaneous frequency) [*F*(1,7.02) = 8.129, *p* = 0.025] of spontaneous excitatory currents (sEPSCs), while there was no change in the amplitude [*F*(2, 42) = 1.763, *p* = 0.184] of sEPSCs when compared to controls (Fig. 1f-i). Measuring spontaneous inhibitory currents (sIPSCs) resulted in a trend of diminished frequency [*F*(2, 9.78) = 2.906, *p* = 0.102] while spatial distribution (instantaneous frequency) [*F*(2, 10.92) = 1.973, *p* = 0.185] and amplitude [*F*(2, 8.41) = 0.458, *p* = 0.458] remained unchanged when compared to controls (Fig. 1f-i). Furthermore, when comparing the mCherry and Saline groups in a post-hoc contrast, no significant differences were observed, signaling the lack of non-ligand dependent effects of the hM4D receptor (EPSC frequency [z(∞) = 0.501, *p*_bonferroni_ = 1.000]; EPSC median instantaneous frequency [z(∞) = 1.097, *p*_bonferroni_ = 0.818]; and no overall effect in EPSC amplitude, IPSC frequency, IPSC instantaneous frequency, or IPSC amplitude. These results indicate that adolescent thalamocortical activity causes significant alterations to excitatory transmission in the superficial regions of the mPFC, consistent with what we previously found in adult mice. (Benoit et al., 2022).

### Adolescent thalamocortical activity regulates excitatory and inhibitory transmission to superficial layer II/III cortico-accumbens projections

We then determined whether the observed effects of adolescent thalamocortical inhibition includes the layer II/III cortico-accumbens cell populations. To this end, we injected the rgGFP virus into the nucleus accumbens (NAc) (Fig. 2a-b) and recorded from the superficial layers (II/III) of the mPFC. We found that in superficial NAc-projecting neurons, excitatory and inhibitory transmission were both dampened following adolescent thalamo-cortical inhibition. For excitatory currents, the frequency [*F*(1, 47) = 9.354, *p* = 0.004] was significantly decreased but not the spatial distribution (instantaneous frequency) [*F*(1, 6.77) = 1.926, *p* =0.208] or the amplitude [*F*(1, 6.19) = 3.313, *p* = 0.117] (Fig. 2e-f). For inhibitory transmission in superficial layers, sIPSC frequency was reduced [*F*(1, 4.89) = 9.451, *p* =0.028] but not the spatial distribution (instantaneous frequency) [*F*(1, 7.79) = 0.955, *p* =0.358] or the amplitude [*F*(1, 7.03) = 3.402, *p* = 0.107] (Fig. 2g-h). These data show that excitatory and inhibitory inputs are decreased onto layer II/III accumbens-projecting neurons.

**Figure 2.**
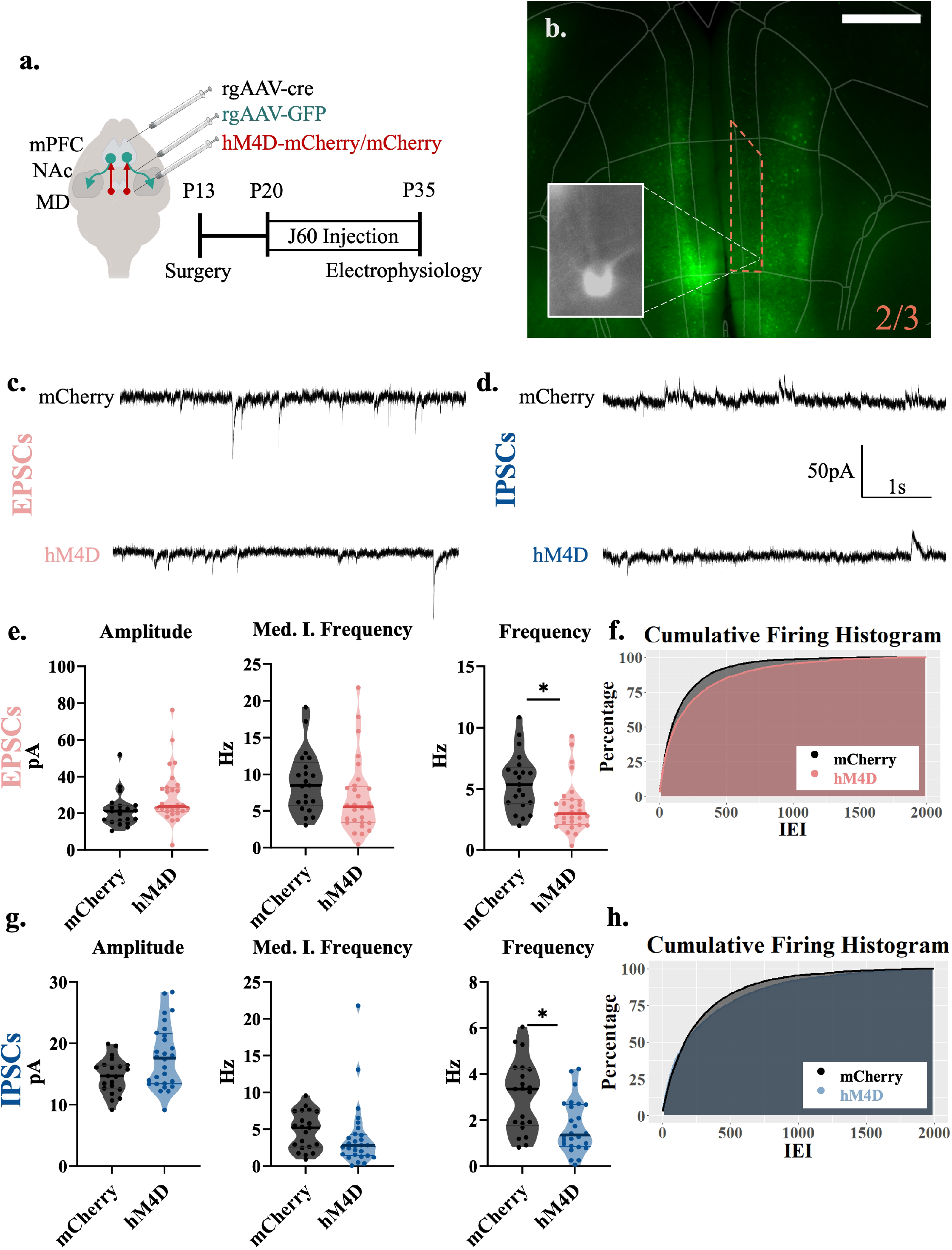
Adolescent thalamocortical activity regulates excitatory and inhibitory transmission to superficial layer II/III cortico-accumbens projections. **a**, Adolescent experimental timeline and schematic. Whole-cell patch clamp recordings were made from accumbens-projecting pyramidal cells in layer II/III of the mPFC from hM4D and control mice; Created with https://biorender.com. **b**, Example image illustrating rgGFP positive, accumbens-projecting cells in the prelimbic subregion of the medial prefrontal cortex, orange dotted outline represents layer II/III. Scale bar represents 500µm. **c**, Representative traces showing sEPSCs of hM4D animals and mCherry controls. **d**, Representative traces showing sIPSCs of hM4D animals and mCherry controls**. e**, sEPSC frequency (right) is significantly reduced in accumbens-projecting cortical cells following adolescent thalamic inhibition relative to control mice, but sEPSC amplitude (left) and median instantaneous frequency (middle) are unchanged; mCherry control, *n* = 21 cells and 5 animals; hM4D, *n* = 28 cells and 6 animals; frequency: mCherry control 5.367(2.717) Hz, hM4D 2.975(1.871) Hz.; median instantaneous frequency: mCherry control 8.514(4.842) Hz and hM4D 5.579(4.803) Hz; amplitude: mCherry control 21.051(7.992) pA and hM4D 23.648(12.495) pA. All reported values are Median(Interquartile Range). **f**, Graph of the cumulative firing distribution for sEPSCs in L II/III accumbens-projecting pyramidal cells in the mPFC, IEI represents the interevent-interval between spontaneous events. **g**, sIPSC frequency (right) is significantly reduced in accumbens-projecting cortical cells following adolescent thalamic inhibition relative to control mice, but sIPSC amplitude (left) and median instantaneous frequency (middle) are unchanged; mCherry control, *n* = 21 cells and 5 animals; hM4D, *n* = 28 cells and 6 animals; frequency: mCherry control 3.350(2.350) Hz, hM4D 1.417(1.762) Hz; median instantaneous frequency: mCherry control 5.195(4.817) Hz and hM4D 2.776(2.753) Hz; amplitude: mCherry control 14.618(3.768) pA and hM4D 17.597(7.919) pA. All reported values are Median(Interquartile Range). **h**, Graph of the cumulative firing distribution for sIPSCs in L II/III accumbens-projecting pyramidal cells in the mPFC. *P<0.05.

### Adolescent thalamocortical activity regulates excitatory and inhibitory transmission to cortico-thalamic projections

We then determined the effects of chronic thalamocortical inhibition on a different subcortical projection, cortical cells from deep layers V/VI projecting to the MD thalamus. A rgGFP virus was injected into the MD and whole cell patch clamp recordings were taken in thalamic-projecting cortical cells (Fig. 3a-b). Like for non-specific layer II/III and cortico-accumbens neurons, we observed significant decreases in excitatory synaptic transmission (Fig. 3e-h). Significant decreases were observed in both the frequency [*F*(1, 6.73) = 12.960, *p* = 0.009] and spatial distribution (instantaneous frequency) [*F*(1, 6.10) = 13.045, *p* = 0.011] of sEPSCs but the amplitude [*F*(1, 5.36) = 0.689, *p* = 0.442] of sEPSCs remained unchanged (Fig. 3e-f). In addition, the frequency [*F*(1, 6.39) = 7.829, *p* = 0.029] and spatial distribution (instantaneous frequency) [*F*(1, 5.96) = 6.501, *p* =0.044] but not the amplitude [*F*(1, 4.77) = 0.991, *p* = 0.367] of sIPSCs was reduced after P20-35 thalamic inhibition (Fig. 3g-h). These data show that, like for superficial layer II/III accumbens projections, adolescent thalamo-prefrontal inhibition also decreases excitatory and inhibitory synaptic inputs to deep layer cortico-thalamic neurons.

**Figure 3.**
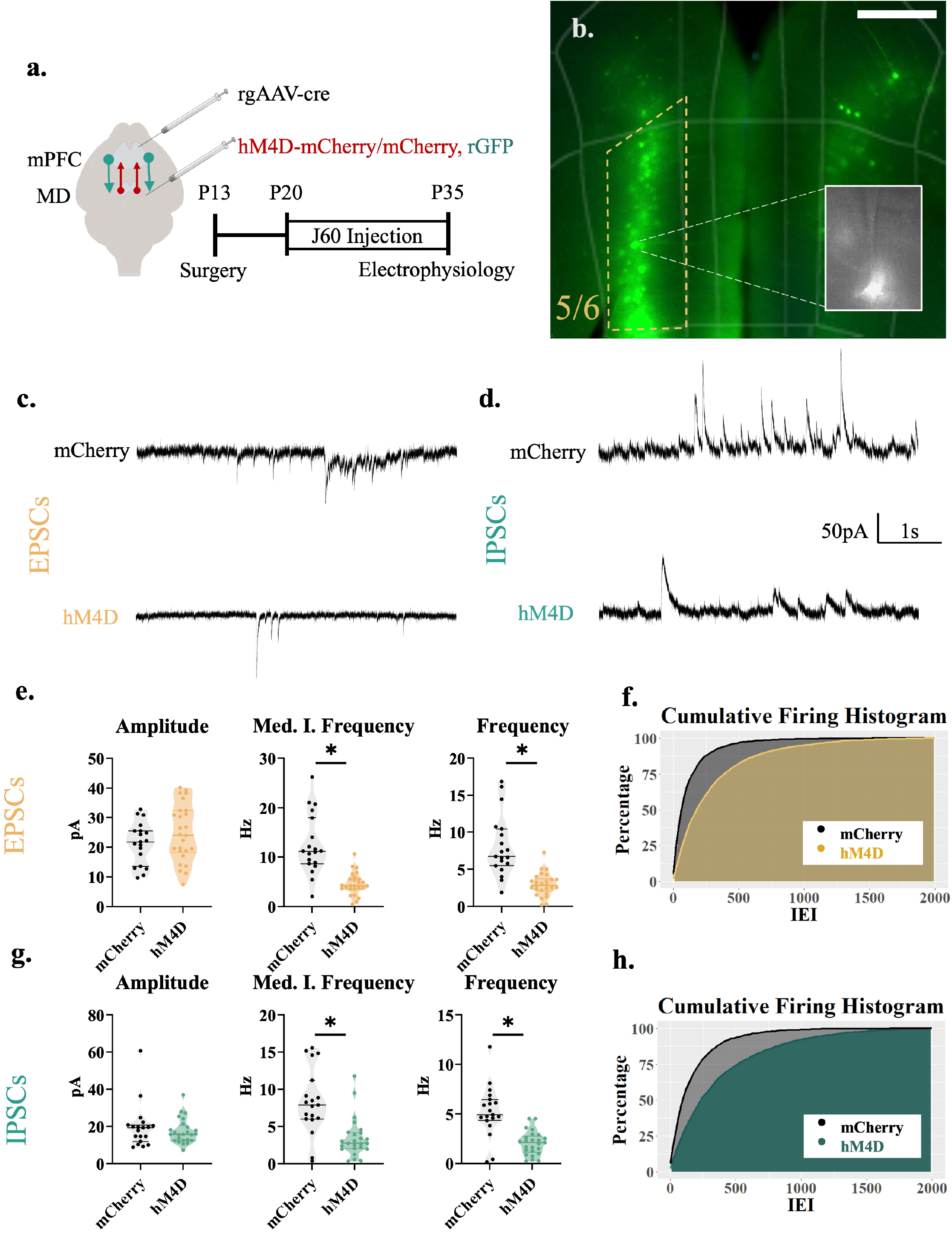
Adolescent thalamocortical activity regulates excitatory and inhibitory transmission to cortico-thalamic projections. **a**, Adolescent experimental timeline, and schematic. Whole-cell patch clamp recordings were made from cortical-projecting pyramidal cells in layer V/VI of the mPFC from hM4D and control mice. These pyramidal cells receive excitatory inputs from the thalamus and superficial layers of the cortex as well as inhibitory inputs from local interneurons; created with https://biorender.com. **b**, Example image illustrating rgGFP positive cells in the prelimbic subregion of the medial prefrontal cortex. All recordings for this experiment were conducted in this region approximately 500-600µm from the pial surface L V/VI (yellow outline). Scale bar represents 300µm **c**, Representative traces showing sEPSCs of hM4D animals and mCherry controls. **d**, Representative traces showing sIPSCs of hM4D animals and mCherry controls**. e**, sEPSC frequency (right) and median instantaneous frequency (middle) is significantly reduced in thalamic-projecting cortical cells following adolescent thalamic inhibition relative to control mice, but sEPSC amplitude (left) is unchanged; mCherry control, *n* = 19 cells and 4 animals; hM4D, *n* = 26 cells and 5 animals; frequency: mCherry control 6.717(4.392) Hz and hM4D 2.833(1.408) Hz; median instantaneous frequency: mCherry control 11.173(7.187) Hz and hM4D 4.204(1.997) Hz; amplitude: mCherry control 21.764(9.897) pA and hM4D 24.043(13.508) pA. All reported values are Median(Interquartile Range). **f**, Graph of the cumulative firing distribution for sEPSCs in L V/VI thalamic-projecting pyramidal cells in the mPFC, IEI represents the interevent-interval between spontaneous events. **g**, sIPSC frequency (right) and median instantaneous frequency (middle) is significantly reduced in thalamic-projecting cortical cells following adolescent thalamic inhibition relative to control mice, but sIPSC amplitude (left) is unchanged; mCherry control, *n* = 19 cells and 4 animals; hM4D, *n* = 26 cells and 5 animals; frequency: mCherry control 4.917(1.825) Hz and hM4D 2.100(1.350) Hz; median instantaneous frequency: mCherry control 7.896(4.100) Hz and hM4D 2.755(2.081) Hz; amplitude: mCherry control 19.267(7.264) pA and hM4D 15.720(7.127) pA. All reported values are Median(Interquartile Range). **h**, Graph of the cumulative firing distribution for sIPSCs in L V/VI thalamic-projecting pyramidal cells in the mPFC. *P<0.05.

### Adolescent thalamocortical activity regulates the amplitude of excitatory and inhibitory currents in deep layer (V/VI) accumbens-projecting cortical cells

Accumbens-projecting prefrontal neurons are located in both superficial and deep cortical layers (Anastasiades and Carter, 2021). We therefore also recorded accumbens-projecting cells in layer V/VI. We found significant increases to excitatory and inhibitory current amplitude, and no changes in frequency or spatial distribution, in deep layer accumbens-projecting cells (Fig 4e-h). When measuring excitatory currents in deep layers, amplitude was significantly increased [*F*(1, 29) = 8.976, *p* =0.006] however, frequency [*F*(1, 5.49) = 7.841×10^-5^, *p* =0.993] and spatial distribution (instantaneous frequency) [*F*(1, 6.24) = 0.196, *p* =0.673] remained unchanged (Fig 4e-f). Similar to excitatory transmission, for inhibitory currents in deeper layers, amplitude was significantly increased [*F*(1, 4.25) = 14.809, *p* =0.013] however, frequency [*F*(1, 6.82) = 0.417, *p* =0.539] and spatial distribution (instantaneous frequency) [*F*(1, 6.53) = 0.910, *p* =0.374] remained unchanged (Fig. 4 g-h). Thus, unlike layer II/III accumbens projection neurons, excitatory and inhibitory drive onto layer V/VI accumbens neurons is not decreased but it is enhanced although at the level of amplitude and not frequency.

**Figure 4.**
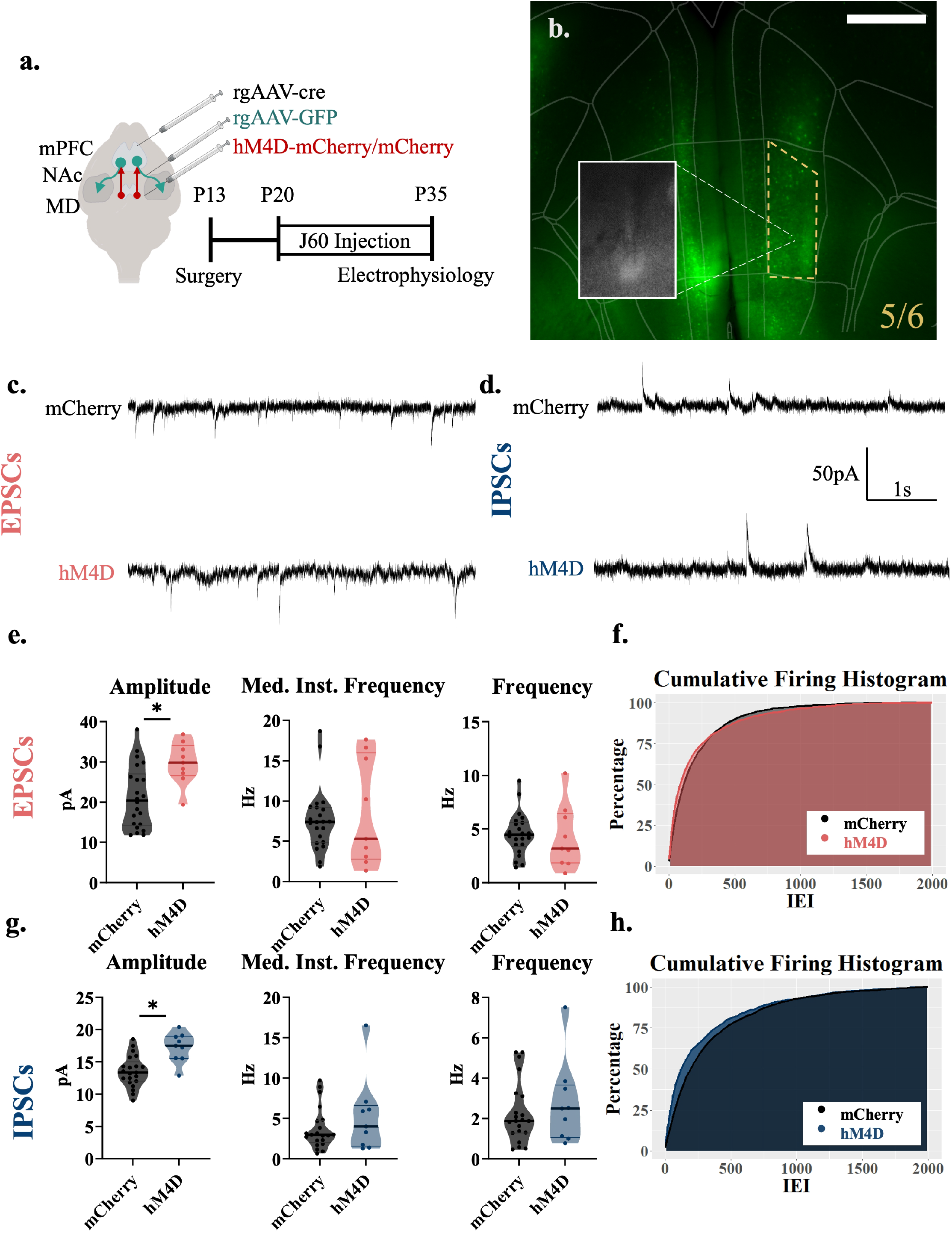
Adolescent thalamocortical activity results in enhanced amplitude in deeper (V/VI) accumbens-projecting cortical cells. **a**, Adolescent experimental timeline and schematic. Whole-cell patch clamp recordings were made from accumbens-projecting pyramidal cells in layer V/VI of the mPFC from hM4D and control mice; Created with https://biorender.com. **b**, Example image illustrating rgGFP positive, accumbens-projecting cells in the prelimbic subregion of the medial prefrontal cortex, Layer V/VI (yellow outline). Scale bar represents 500µm. **c**, Representative traces showing sEPSCs of hM4D animals and mCherry controls. **d**, Representative traces showing sIPSCs of hM4D animals and mCherry controls**. e**, sEPSC amplitude (left) is significantly increased in accumbens-projecting cortical cells following adolescent thalamic inhibition relative to control mice, but sEPSC frequency (right) and median instantaneous frequency (middle) are unchanged; mCherry control, *n* = 22 cells and 5 animals; hM4D, *n* = 9 cells and 4 animals; frequency: mCherry control 4.450(2.012) Hz, hM4D 3.167(4.217) Hz.; median instantaneous frequency: mCherry control 7.371(4.043) Hz and hM4D 5.290(12.190) Hz; amplitude: mCherry control 20.460(11.638) pA and hM4D 29.858(5.869). All reported values are Median(Interquartile Range). **f**, Graph of the cumulative firing distribution for sEPSCs in L II/III accumbens-projecting pyramidal cells in the mPFC, IEI represents the interevent-interval between spontaneous events. **g**, sIPSC frequency (right) is significantly reduced in accumbens-projecting cortical cells following adolescent thalamic inhibition relative to control mice, but sIPSC amplitude (left) and median instantaneous frequency (middle) are unchanged; mCherry control, *n* = 21 cells and 5 animals; hM4D, *n* = 28 cells and 6 animals; frequency: mCherry control 1.883(1.817) Hz, hM4D 2.50(2.350) Hz; median instantaneous frequency: mCherry control 2.997(2.405) Hz and hM4D 4.005(4.349) Hz; amplitude: mCherry control 13.362(2.306) pA and hM4D 17.495(3.331). All reported values are Median(Interquartile Range). **h**, Graph of the cumulative firing distribution for sIPSCs in L V/VI accumbens-projecting pyramidal cells in the mPFC. *P<0.05.

### Adolescent thalamocortical activity regulates cross-hemispheric inhibition

To determine how adolescent thalamocortical inhibition impacts cortico-cortical circuitry, a rgGFP virus was injected into the dextral prelimbic mPFC and whole cell patch clamp recordings were taken in callosally-projecting cortical cells in the sinistral hemisphere (Fig. 5a-b). No significant differences were observed in the frequency [*F*(1, 7.41) = 0.012, *p* = 0.917], spatial distribution (instantaneous frequency) [*F*(1, 7.30) = 0.399, *p* = 0.547] or amplitude) [*F*(1, 6.37) = 2.865, *p* = 0.134] of excitatory events in cortical-projecting cortical cells (Fig. 5e-f). Unlike excitatory transmission, inhibitory transmission significantly increased in the distribution of events (instantaneous frequency) [*F*(1, 5.95) = 8.023, *p* = 0.030] and trended in frequency [*F*(1, 7.32) = 4.185, *p* = 0.078] but not amplitude [*F*(1, 62) = 0.112, *p* = 0.912] after adolescent thalamocortical inhibition (Fig. 5g-h). These findings were unexpected since callosally-projecting prelimbic neurons are enriched in layer II/III (Anastasiades et al., 2018) and the non-selective analysis of layer II/III neurons displayed a decrease in the frequency of excitatory currents and no increase in inhibition.

**Figure 5.**
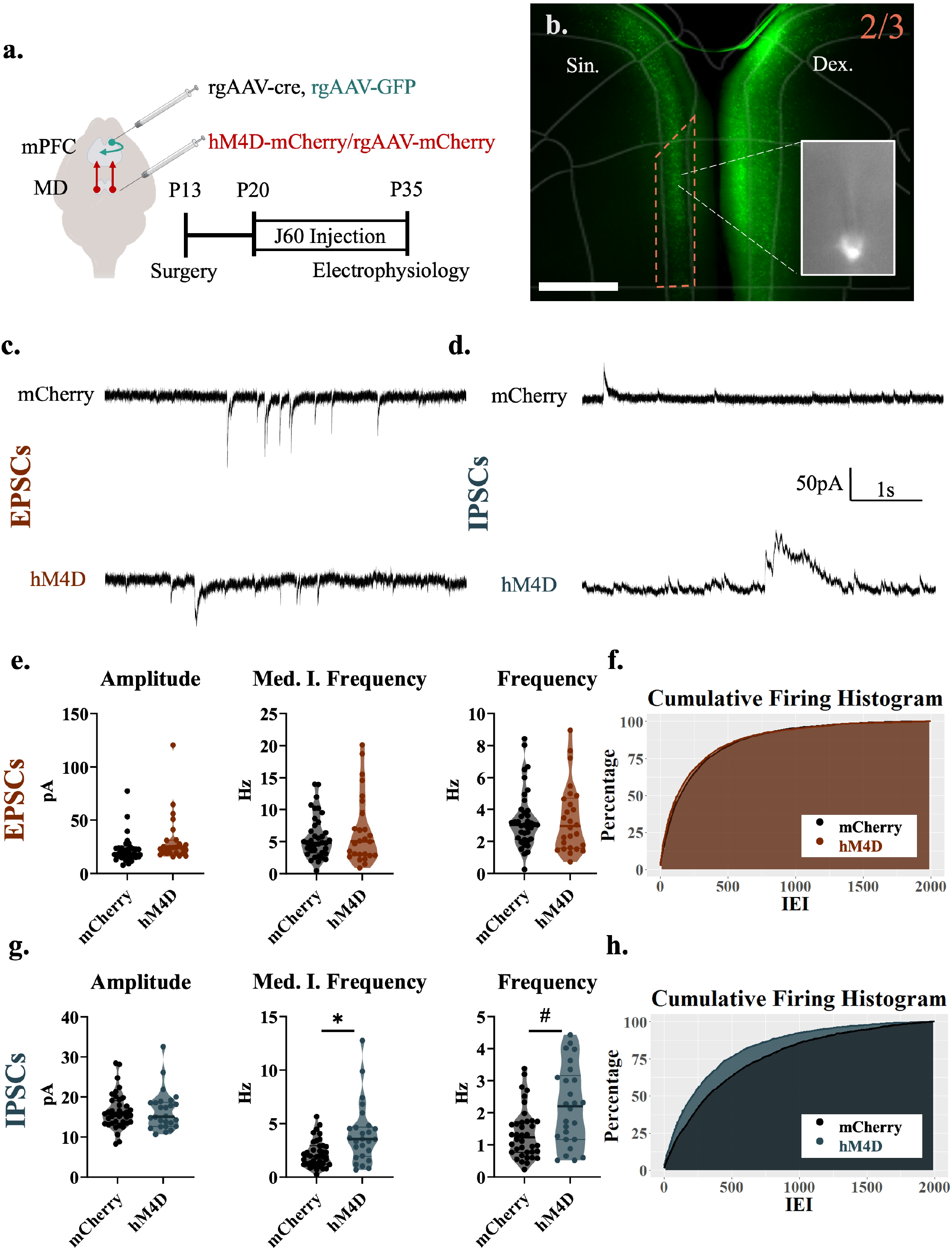
Adolescent thalamocortical activity regulates cross-hemispheric inhibition. **a**, Adolescent experimental timeline and schematic. Whole-cell patch clamp recordings were made from cortical-projecting pyramidal cells in layer II/III of the mPFC from hM4D and control mice; Created with https://biorender.com. **b**, Example image illustrating rgGFP positive, cortical-projecting cells in the prelimbic subregion of the medial prefrontal cortex (orange outline). Virus was injected into the dextral (right) hemisphere and cells were recorded only in the sinistral (left) hemisphere. Scale bar represents 500µm. **c**, Representative traces showing sEPSCs of hM4D animals and mCherry controls. **d**, Representative traces showing sIPSCs of hM4D animals and mCherry controls**. e**, sEPSC frequency (right), sEPSC amplitude (left) and sEPSC median instantaneous frequency are unchanged in cortical-projecting cortical cells following adolescent thalamic inhibition relative to control mice; mCherry control, *n* = 38 cells and 5 animals; hM4D, *n* = 26 cells and 5 animals; frequency: mCherry control 3.042(1.713) Hz and hM4D 2.95(2.933) Hz; median instantaneous frequency: mCherry control 4.807(3.944) Hz and hM4D 5.042(5.890) Hz; amplitude: mCherry control 17.876(7.549) pA and hM4D 23.951(8.397) pA. All reported values are Median(Interquartile Range). **f**, Graph of the cumulative firing distribution for sEPSCs in cortical-projecting pyramidal cells in the mPFC, IEI represents the interevent-interval between spontaneous events. **g**, sIPSC median instantaneous frequency (middle) is significantly increased and frequency (right) is trending towards an increase in cortical-projecting cortical cells following adolescent thalamic inhibition relative to control mice, while amplitude (left) is unchanged; mCherry control, *n* = 38 cells and 5 animals; hM4D, *n* = 26 cells and 5 animals; frequency: mCherry control 1.233(0.954) Hz and hM4D 2.208(1.887) Hz; median instantaneous frequency: mCherry control 1.937(1.676) Hz and hM4D 5.551(2.498); amplitude: mCherry control 15.735(5.268) pA and hM4D 15.099(5.667) pA. All reported values are Median(Interquartile Range). **h**, Graph of the cumulative firing distribution for sIPSCs in cortical-projecting pyramidal cells in the mPFC. *P<0.05, #P<0.10.

### Excitation-Inhibition balance is only altered in callosally-projecting mPFC neurons

Excitation and inhibition are well calibrated in cortical neurons (Okun and Lampl, 2008) and in agreement with this we found a strong correlation in the frequency of excitatory and inhibitory currents for non-specific layer II/III cortical cells [R^2^=0.524, P<0.001], cortico-accumbens cells [R^2^=0.811, P<0.001], cortico-thalamic [R^2^=0.850, P<0.001], and cortico-cortical cells [R^2^=0.550, P<0.001]. In many cell populations, we observed changes in both excitation and inhibition, suggesting homeostatic mechanisms may compensate for reduced excitatory drive from the thalamus. We therefore calculated the excitation inhibition-balance in the different analyzed population of neurons. No changes in the excitation-inhibition balance were measured in layer II/III and V/VI cortico-accumbens-projecting neurons, and cortico-thalamic neurons (Fig. 6b-e). In contrast, when analyzing the ratio of excitatory to inhibitory currents in contralaterally-projecting mPFC neurons, the frequency [*F*(1, 11.37) = 5.697, *p* = 0.035] and spatial distribution (instantaneous frequency) [*F*(1, 9.34) = 5.744, *p* = 0.039] of excitatory to inhibitory currents was significantly weighted towards inhibition (Fig. 6f).

**Figure 6.**
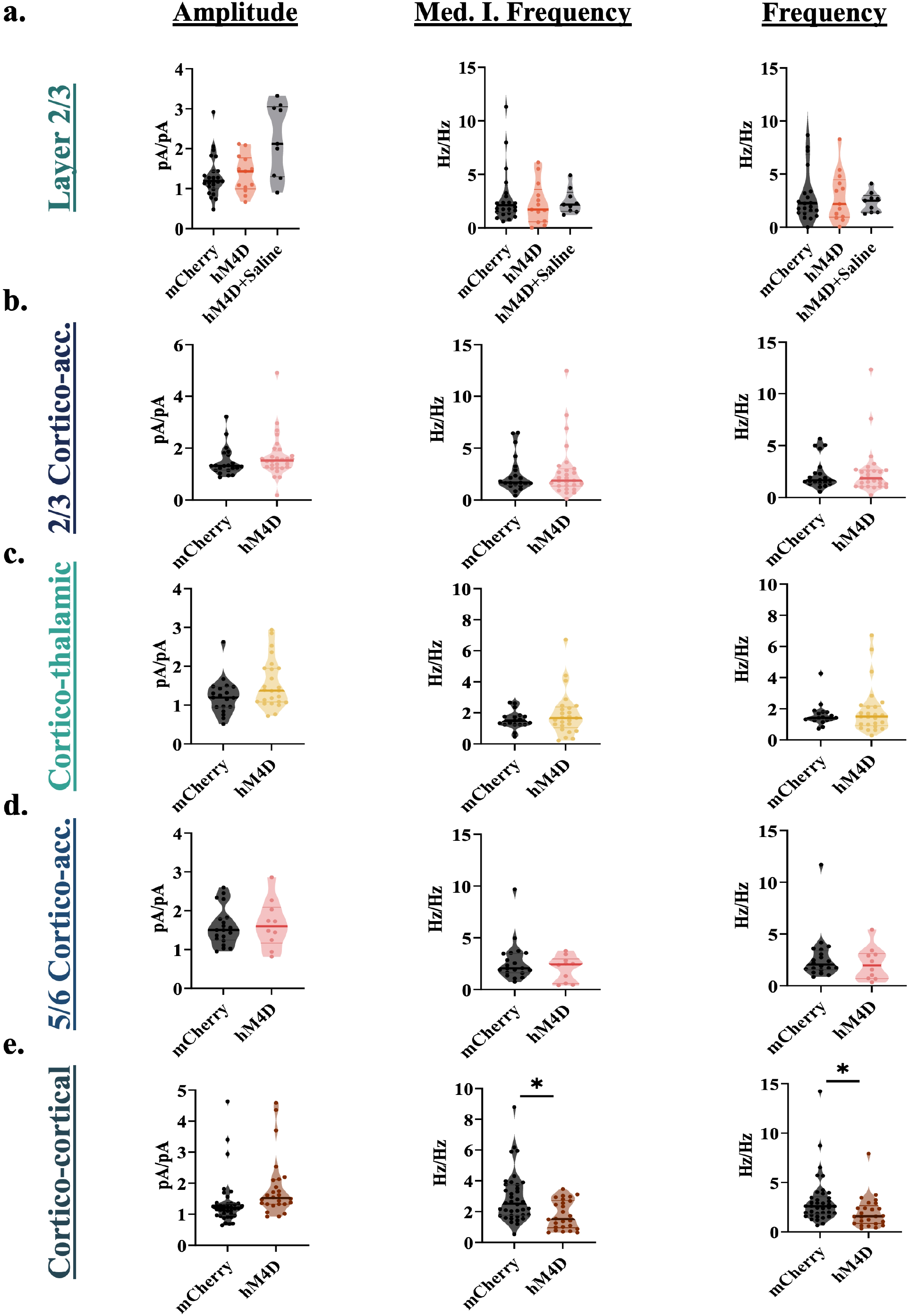
Adolescent thalamocortical inhibition disrupts E/I Ratio in cortical-projecting cortical cells. **a,** Excitatory/Inhibitory ratio in nonspecific layer II/III cells for amplitude (left), median instantaneous frequency (middle), and frequency (right). There was no significant difference in E/I ratio between mCherry 1.188(0.400) pA/pA, hM4D 1.432(0.704) pA/pA or hM4D+Saline 2.005(1.247) pA/pA for amplitude [*f* = 2.567, d.f. = 2, 11.07 *P* = 0.121]. There was no significant difference in E/I ratio between mCherry 2.131(2.020) Hz/Hz, hM4D 1.736(2.409) Hz/Hz or hM4D+Saline 2.183(1.306) Hz/Hz for median instantaneous frequency [*f* = 0.713, d.f. = 2, 11.65 *P* = 0.511]. There was no significant difference in E/I ratio between mCherry 2.265(4.677) Hz/Hz, hM4D 2.186(3.134) Hz/Hz or hM4D+Saline 2.490(1.325) Hz/Hz for median instantaneous frequency [*f* = 1.209, d.f. = 2, 43 *P* = 0.308]. **b,** Excitatory/Inhibitory ratio in layer II/III accumbens-projecting cortical cells for amplitude (left), median instantaneous frequency (middle), and frequency (right). There was no significant difference in E/I ratio between mCherry 1.316(0.587) pA/pA or hM4D 1.524(0.505) pA/pA for amplitude [*f* = 0.543, d.f. = 1, 6.59 *P* = 0.487]. There was no significant difference in E/I ratio between mCherry 1.686(1.368) Hz/Hz and hM4D 1.999(1.715) Hz/Hz for median instantaneous frequency [*f* = 0.836, d.f. = 1, 7.91 *P* = 0.388]. There was no significant difference in E/I ratio between mCherry 1.663(0.963) Hz/Hz and hM4D 1.865(1.406) Hz/Hz for frequency [*f* = 0.750, d.f. = 1, 8.10 *P* = 0.411]. **c,** Excitatory/Inhibitory ratio in layer V/VI thalamic-projecting cortical cells for amplitude (left), median instantaneous frequency (middle), and frequency (right). There was no significant difference in E/I ratio between mCherry 1.195(0.500) pA/pA or hM4D 1.376(0.858) pA/pA for amplitude [*f* = 3.625, d.f. = 1, 42 *P* = 0.064]. There was no significant difference in E/I ratio between mCherry 1.414(0.485) Hz/Hz and hM4D 1.660(1.143) Hz/Hz for median instantaneous frequency [*f* = 0.975, d.f. = 1, 5.21 *P* = 0.367]. There was no significant difference in E/I ratio between mCherry 1.414(0.485) Hz/Hz and hM4D 1.503(1.136) Hz/Hz for frequency [*f* = 1.115, d.f. = 1, 5.31 *P* = 0.337]. **d,** Excitatory/Inhibitory ratio in layer V/VI accumbens-projecting cortical cells for amplitude (left), median instantaneous frequency (middle), and frequency (right). There was no significant difference in E/I ratio between mCherry 1.467(0.624) pA/pA or hM4D 1.730(0.618) pA/pA for amplitude [*f* = 2.220, d.f. = 1, 28 *P* = 0.147]. There was no significant difference in E/I ratio between mCherry 2.060(1.840) Hz/Hz and hM4D 2.432(1.435) Hz/Hz for median instantaneous frequency [*f* = 0.412, d.f. = 1, 12.33 *P* = 0.533]. There was no significant difference in E/I ratio between mCherry 1.829(1.623) Hz/Hz and hM4D 2.658(1.765) Hz/Hz for frequency [*f* = 0.014, d.f. = 1, 12.10 *P* = 0.908]. **e,** Excitatory/Inhibitory ratio in cortical-projecting cortical cells for amplitude (left), median instantaneous frequency (middle), and frequency (right). There was no significant difference in E/I ratio between mCherry 1.209(0.359) pA/pA or hM4D 1.473(0.403) pA/pA for amplitude [*f* = 1.099, d.f. = 1, 59 *P* = 0.299]. There was a significant decrease in E/I ratio between mCherry 2.513(1.915) Hz/Hz and hM4D 1.517(1.670) Hz/Hz for median instantaneous frequency. There was also a significant decrease in E/I ratio between mCherry 2.594(1.607) Hz/Hz and hM4D 1.564(1.544) Hz/Hz for frequency. All reported values are Median(Interquartile Range). *P<0.05.

### P20-P35 thalamocortical inhibition produces cognitive deficits in adult mice

We then determined whether thalamocortical inhibition from P20-P35 has similar long-lasting effects on cognition compared to P20-50 inhibition. J60 was administered from P20-P35 to mice expressing hM4D or a control fluorophore (GFP) in thalamus-prelimbic cortex-projecting neurons. Behavioral testing was conducted 55 days later, at P90 (Fig. 7a). After thalamocortical inhibition, at P90, mice performed the attentional set-shifting task (ASST). For initial acquisition, mice were trained to associate a cinnamon scent with a reward irrespective of bedding medium. After meeting criterion, an extra-dimensional set shift (EDSS) occurred, and mice had to associate the paper bedding with a reward irrespective of scent (Fig. 7d). Adolescent thalamo-prefrontal inhibition did not affect the initial acquisition of the task [t=0.9647, df=22, p=0.3452] but delayed acquisition of the extra-dimensional set shift compared to controls [t=2.442, df=22, *p=0.0354] (Figure 7e). Adolescent-inhibited hM4D animals committed more perseverative errors than control animals [t=2.461, df=22, *p=0.0222] (Figure 7f). Perseverative errors are when the mouse continues with the same response strategy following a set shift. However, adolescent-inhibited hM4D animals also committed more random errors than control animals and the deficit was not due to a specific error type [t=2.786, df=22, *p=0.0108] (Figure 7f). These data indicate that a restricted critical window (P20-P35) is sufficient to manifest deficits observed with a longer period of inhibition (P20-P50) (Benoit et al., 2022).

**Figure 7.**
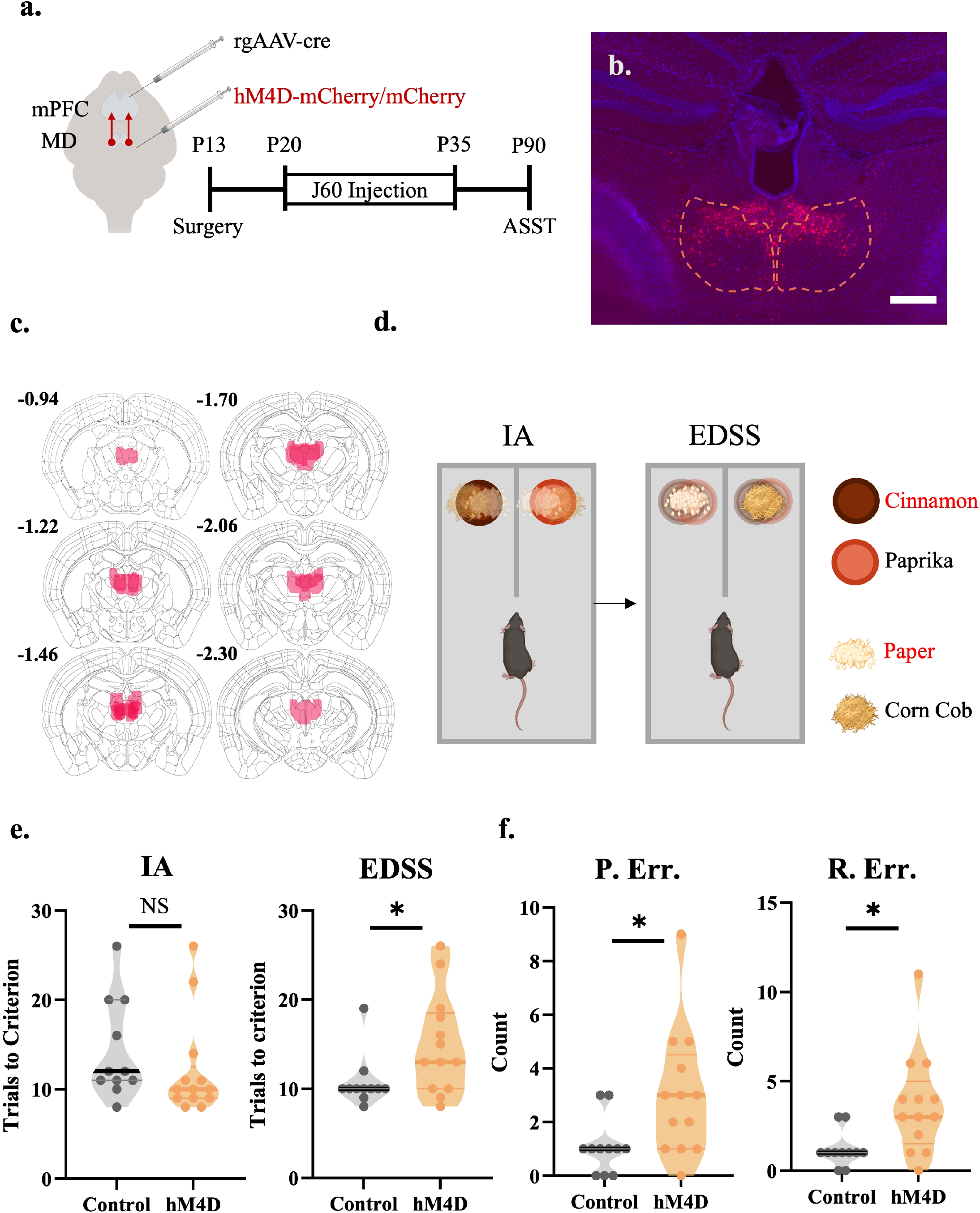
P20-P35 thalamocortical inhibition produces cognitive deficits in adult mice. **a,** Adolescent experimental timeline, and schematic. J60 was administered from P20-P35 to mice expressing hM4D or a control fluorophore (GFP) in the thalamus. Behavioral testing was conducted 55 days later, at P90. Created with https://biorender.com **b,** Representative image of hM4D expression in the MD of adult mice. Scale bar represents 300µm. **c,** Representative viral spread of hM4D receptor in mice that underwent the ASST task. **d,** Schematic of the attentional set-shifting task (ASST). Mice were trained to associate a cinnamon scent with a reward irrespective of bedding medium (IA). After meeting criterion, an extra-dimensional set shift occurred (EDSS) and mice had to associate the paper bedding with a reward irrespective of scent. Red is representative of the rewarded stimulus. **e,** Adolescent-inhibited hM4D animals are no different than controls in the initial acquisition (IA) of the ASST (left, Control: n=11 animals, 14.27± 2.276 trials, hM4D: n=13 animals, hM4D: 12.08±2.276 trials; two-sided unpaired t-test, t=0.9647, df=22, p=0.3452) but take significantly more trials in the extra-dimensional set shift (EDSS) than controls (right, Control: n=11 animals, 10.73±1.871 trials, hM4D: n=13 animals, hM4D: 14.92±1.871 trials; unpaired t-test, t=2.442, df=22, *p=0.0354). **f,** Adolescent-inhibited hM4D animals committed more perseverative errors (P. Err.) than control animals (left, Control: n=11 animals, 1.091±0.7759 trials, hM4D: n=13 animals, hM4D: 3.000±0.7759 trials; unpaired t-test, t=2.461, df=22, *p=0.0222). Perseverative errors are when the mouse continues with the same response strategy following a rule switch. Adolescent-inhibited hM4D animals committed more random errors (P. Err.) than control animals (right, Control: n=11 animals, 1.182±0.9010 trials, hM4D: n=13 animals, hM4D: 3.692±0.9010 trials; unpaired t-test, t=2.786, df=22, *p=0.0108) non-perseverative errors are generally considered to be random.

## Discussion

To determine thalamocortical activity’s importance for prefrontal circuit maturation we inhibited thalamo-mPFC projections from P20-35 and recorded from prelimbic neurons twenty-four hours later. We made several observations. First, our findings reveal that excitatory transmission to layer-II/III is already decreased during adolescence, suggesting the adult excitation loss (Benoit et al., 2022) does not require maturation to adulthood but arises during adolescence. Second, we measured decreased excitatory and inhibitory transmission to layer-II/III NAc- and layer-V/VI thalamic-projecting neurons. These inhibitory changes suggest adolescent-thalamocortical-inhibition leads to alterations in prefrontal connectivity beyond the loss of excitatory thalamic input, previously observed at the anatomical level at P35 (Benoit et al., 2022). Third, we observed a surprising increase in excitatory and inhibitory transmission to layer-V/VI NAc-projecting neurons in amplitude but not frequency, suggesting a post-synaptic adaptation.

While the specificity of this effect is unclear, NAc-projecting cells in deep layers have distinct molecular characteristics from superficial layers, which may differentially impact their response to adolescent-thalamocortical-inhibition (Bhattacherjee et al., 2019). Fourth, we measured enhanced inhibition in cortico-cortical connectivity, providing further evidence for a complex adaptation in response to adolescent-thalamocortical-inhibition. Intriguingly, while excitatory and inhibitory (E/I) drive was altered onto subcortical projection neurons, a change in the E/I balance was only measured in cross-hemispheric mPFC projections and not in the three subcortical projections analyzed. As functional stability in neural circuits can be sustained by homeostatic re-balancing of E/I inputs, the largest functional effect of adolescent-thalamocortical-inhibition on prefrontal circuitry may be a decrease in the activity of callosally-projecting mPFC pyramidal cells. While difficult to predict neuronal activity in-vivo from slice physiology, enhanced inhibition, and less activation of callosal-projecting neurons during adolescence might have long-lasting consequences for mPFC function. Finally, we established that this narrowed window of inhibition (P20-P35) is sufficient to produce cognitive deficits in the adult mouse.

### Decreased excitatory and inhibitory transmission in mPFC arises during adolescence

Previous work demonstrated that adolescent-thalamocortical-inhibition (P20-P50) resulted in cognitive and physiological deficits in the adult mouse, including a loss of excitatory transmission in non-projection-specific layer-II/III pyramidal cells. Here, we observe deficits are established during adolescence, prior to being sustained into adulthood. This excitatory dampening onto layer-II/III neurons could result from the loss of MD innervation to pyramidal cells and/or a loss of local excitation. The bimodal distribution of the effects in layer-II/III led us to hypothesize that adolescent thalamocortical-inhibition may be differentially affecting subpopulations of neurons in the subregion.

Excitatory and inhibitory currents were both decreased in NAc-projecting pyramidal cells following adolescent-thalamocortical-inhibition. These data show that adolescent-thalamocortical-inhibition may have widespread effects on diverse cell populations, including mPFC-NAc projections that are essential for reward driven motivated behaviors (Kalivas et al., 2005; Mannella et al., 2013; McGlinchey et al., 2016).

The decrease in inhibitory transmission could be due to a reduction in thalamic projections onto interneurons (Yang et al., 2021). The MD directly innervates layer-II/III parvalbumin (PV+) and other interneurons, providing a source of feedforward inhibition to cortical projection neurons (Delevich et al., 2015; Canetta et al., 2020). Changes in interneuron-driven inhibition in NAc-projecting cells measured in slice could reflect either reduced excitatory drive onto PV+ cells, or adaptations within PV+ interneurons following thalamic inhibition. To determine whether adolescent-thalamocortical-inhibition also alters excitatory and inhibitory connections in deeper layers of the mPFC, we next recorded from cortico-thalamic neurons in layer-V/VI.

We found that adolescent thalamocortical-inhibition resulted in a decrease in the frequency of excitatory and inhibitory currents in thalamus-projecting cells in layer-V/VI of the mPFC, like the effects seen in superficial NAc-projecting cells. The MD primarily projects to layer-II/III; however, slice physiology has demonstrated monosynaptic, direct-reciprocal connection to the deeper layers (V/VI) of the cortex (Collins et al., 2018). The origins for the excitatory and inhibitory decrease may therefore be like the ones measured in layer-II/III.

### Deeper accumbens-projecting cells show post-synaptic adaptations to MD inhibition

Surprisingly, we found that following adolescent-thalamocortical-inhibition, the amplitude of excitatory and inhibitory currents increased in layer-V/VI accumbens-projecting neurons, while the frequency and spatial distribution remained the same. Changes in amplitude are thought to reflect post-synaptic alterations, this amplitude increase may therefore be a post-synaptic adaptation. It is unclear; however, for what the increased amplitude would be compensating. Although the thalamus makes direct connections onto deep layer pyramidal cells (Collins et al., 2018), it is unknown whether the MD directly innervates deep NAc-projecting cells.

### Adolescent-thalamocortical-inhibition increases inhibitory inputs to callosally-projecting neurons

The frequency of excitatory inputs to callosally-projecting mPFC neurons did not change following adolescent-thalamocortical-inhibition; however, inhibitory inputs were increased. The absence of excitatory transmission changes in this population is surprising, given they receive strong MD inputs (Anastasiades et al., 2018). Why excitatory inputs to this population are spared remains unknown, but it suggests that excitatory inputs from other brain regions may offset the reduction in excitatory inputs from the thalamus. The increase in inhibitory transmission is also surprising given that inhibition was decreased in NAc-projecting cells found in the same cortical layer. Moreover, while changes in E/I, driven by adolescent-thalamocortical-inhibition largely scaled in all other populations, cortico-cortical-projecting cells showed a significant reduction in their sE/IPSC ratio. Overall, E/I ratio stability following adolescent thalamocortical-inhibition displays homeostatic regulation in subcortically projecting neurons as inhibition may be an adaptive consequence to the decrease in excitatory thalamic inputs. However, in cortically projecting neurons, adolescent thalamocortical-inhibition resulted in an inhibitory increase with no effect on excitation, and a significant reduction in E/I balance in these cells. Thus, callosally-projecting neurons may not undergo homeostatic adaptation compared to subcortically-projecting cells. Work in the somatosensory cortex shows subcortically-projecting neurons in layer-V display faster synaptic depression and homeostatic rebound compared to cortically-projecting layer-V cells (Greenhill et al., 2015). If conserved in layer-II/III of the mPFC, this could be a rationale for delayed homeostatic adaptation in cortically-projecting cortical cells. Therefore, we predict cortically-projecting neurons might experience the greatest change following adolescent-thalamocortical-inhibition, with the net effect of being less active in vivo. This change may in turn contribute to a loss of cross-hemispheric cortical communication, hypothesized to be important for cognitive processes (Hasegawa et al., 1998; Holler-Wallscheid et al., 2017). These effects may depress cortical output to other neural circuits, having widespread effects on prefrontal network function.

Alternatively, the increase in callosal inhibition following adolescent-thalamocortical-inhibition may have resulted from interneuron-mediated disinhibition via somatostatin (SST+), vasoactive intestinal polypeptide (VIP+), and PV+ interneurons. Both VIP+ and SST+ interneurons receive MD input and inhibit other interneurons including PV+ interneurons (Ahrlund-Richter et al., 2019; Sun et al., 2019; Canetta et al., 2020). Therefore, the overall increase in inhibition could be due to complex disinhibition between VIP+, SOM+, PV+ and pyramidal cell types. It is unclear why it should be specific only to callosal, intracortical connectivity but some interneurons target callosal projecting pyramidal cells (Pi et al., 2013; Cho et al., 2023).

### Possible mechanisms of mPFC plasticity induced by adolescent-thalamocortical-inhibition

Recent studies also point to intra-cortical mechanisms mediating maturation of cortical circuitry. Anterior cingulate (ACC) neurons show greater excitatory synaptic inputs during adolescence than adulthood and inhibition of ACC to visual cortex projection neurons during adolescence disrupts the maintenance of local connectivity within the ACC (Nabel et al., 2020). By analogy, if adolescent-thalamocortical-inhibition decreases activity of mPFC-thalamic or accumbens projection neurons during adolescence, this may lead to a disruption in the maintenance of local excitation in the mPFC. A different mechanism has been described within the visual cortex. Here, inhibition of layer-II/III neurons during the critical period of primary visual cortex development (P24-29) led to excitatory synaptic scaling and increased intrinsic excitability suggesting homeostatic plasticity as a mechanism affecting visual cortex maturation (Wen and Turrigiano, 2021). Further studies on synaptic scaling could identify the plasticity mechanisms in mPFC circuitry induced by adolescent-thalamocortical-inhibition.

### A narrowed window of adolescent-thalamocortical-inhibition regulates adult cognition

Adolescent-thalamocortical-inhibition from P20-35 elicited similar deficits in attentional set shifting the adult mouse at P90 window compared to a broader time window from P20-50 (Benoit et al., 2022). This narrows the sensitive window, implicating early adolescence as a critical period during which thalamic input regulates prefrontal cortex development.

## Conclusions

In sum, our data demonstrate adolescent-thalamocortical-inhibition results in projection specific changes in synaptic connectivity in the mPFC that become evident already during adolescence. The dysregulation of multiple neural circuits demonstrates a widespread effect of adolescent-thalamocortical-inhibition that impacts more than just thalamocortical circuitry and deficits cannot be explained solely by a lack of anatomical input. Rather, they generate widespread adaptative and possibly maladaptive physiological responses within prefrontal circuitry.

## Future Directions

Future studies directly measuring excitatory and inhibitory currents and using optogenetic approaches to probe functional connectivity in interneurons and pyramidal cells will be able to address how these complex adaptations arise. Studies exploring how these physiological changes will scale into adulthood will be of interest. If they do, it is likely adolescent-thalamocortical-inhibition would lead to changes in prefrontal circuitry that are ‘hardwired’ into adulthood. Alternatively, changes identified in adolescence may be transient, setting in motion altered adult prefrontal function. Finally, future research will address whether disinhibition of callosal-cortical-projecting cells during adolescence following adolescent-thalamocortical-inhibition will rescue long-term negative effects and determine whether adolescence is a potential target for therapeutic intervention.

## Acknowledgments

This work was supported by NIMH R01MH128293 to C.K. and S.C and NIMH R01MH1282771 to S.C.

